# Enzyme kinetics of tobacco Rubisco expressed in *Escherichia coli* varies depending on the small subunit composition

**DOI:** 10.1101/562223

**Authors:** Myat T. Lin, William D. Stone, Vishal Chaudhari, Maureen R. Hanson

## Abstract

Rubisco catalyzes the first step in carbon fixation and has been a strategic target to improve photosynthetic efficiency. In plants, Rubisco is a complex made up of eight large subunits encoded by a chloroplast gene, *rbcL*, and eight small subunits expressed from a nuclear gene family and targeted to chloroplast stroma. Biogenesis of Rubisco in plants requires a chaperonin system composed of Cpn60α, Cpn60β and Cpn20, which helps fold the large subunit, and multiple chaperones including RbcX, Raf1, Raf2 and BSD2, which help the dimerization of the folded large subunits and subsequent assembly with the small subunits into L_8_S_8_ holoenzymes. A recent study successfully assembled functional Arabidopsis Rubisco in *Escherichia coli* by co-expressing the two subunits with Arabidopsis chaperonins and chaperones (Aigner et al., 2017). In this study, we modified the expression vectors used in that study and adapted them to express tobacco Rubisco by replacing the Arabidopsis genes with tobacco ones. Next, we surveyed the small subunits present in tobacco, co-expressed each with the large subunit and successfully produced active tobacco enzymes composed of different small subunits in *E. coli*. These enzymes produced in *E. coli* have carboxylation kinetics very similar to that of the native tobacco Rubisco. We also produced tobacco Rubisco with a recently discovered trichome small subunit in *E. coli* and found that it has a higher catalytic rate and a lower CO_2_ affinity compared to the enzymes with other small subunits. Our improvements in the *E. coli* Rubisco expression system will allow us to probe features of both the chloroplast and nuclear-encoded subunits of Rubisco that affect its catalytic rate and CO_2_ specificity.

## Introduction

In photosynthetic organisms including plants, ribulose-1,5-bisphosphate carboxylase/oxygenase (Rubisco, EC 4.1.1.39) catalyzes the fixation of CO_2_ from air to ribulose-1,5-bisphosphate (RuBP). Thus, Rubisco represents a major gateway for inorganic carbon to enter biosphere and has far-reaching impacts on a global scale. The carboxylation of RuBP by Rubisco is a well-known bottleneck in C_3_ photosynthesis because of its slow catalytic rates with approximately 3 s^−1^ turnover number (*k_cat_*) (Whitney et al., 2011a). Rubisco also catalyzes a competing RuBP oxygenation, generating a toxic byproduct, 2-phosphoglycolate, which has to be recycled through the energy-intensive photorespiration pathway (Bauwe et al., 2012). The efficiency loss due to the RuBP oxygenation depends on temperature, ambient CO_2_ and other environmental conditions and has been estimated to lower the yield of C_3_ crops by about 20-36% (Walker et al., 2016).

An important kinetic parameter of Rubisco is its CO_2_/O_2_ specificity factor (*S*_C/O_), which is the ratio of the catalytic efficiencies of its carboxylation reaction 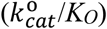 and oxygenation reaction 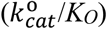, where *k_cat_* and *K* are the catalytic rate and Michaelis-Menten constant respectively (Jordan and Ogren, 1981). Due to mechanistic constraint, a Rubisco that binds its carboxylated intermediates tighter has a higher *S_C/O_*, a lower *K_C_* and a lower 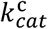 leading to a strong direct relationship between 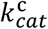 and *K_C_* as well as an inverse relationship between 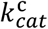 and *S_C/O_* (Tcherkez et al., 2006; Savir et al., 2010). Due to the lack of a CO_2_-concentrating mechanism, C_3_ Rubisco enzymes typically have higher *S_C/O_* and lower 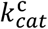 than C_4_ enzymes. To overcome the low catalytic rates, C_3_ plants have evolved to produce much more Rubisco and invest as much as 25% of leaf nitrogen in just one enzyme (Evans, 1989).

The plant enzyme belongs to form I Rubisco and is composed of eight large subunits encoded by a single chloroplast gene and eight small subunits expressed from a family of nuclear genes. In contrast, dinoflagellates, many proteobacteria and archaea produce forms II and III Rubisco without small subunits (Tabita et al., 2008). The active site is formed at the dimeric interface between two large subunits and well conserved among all Rubisco forms. Cyanobacteria and many autotropic prokaryotes also possess form I Rubisco, which has the same subunit composition as in the plant L_8_S_8_ holoenzyme. The large subunits are prone to aggregation and require GroEL/ES-type chaperonin for proper folding followed by assistance from additional chaperones for dimerization and, in the case of form I Rubisco, subsequent assembly with small subunits to form the L_8_S_8_ holoenzyme (Wilson and Hayer-Hartl, 2018). Form I Rubisco from many cyanobacteria can be assembled in *E. coli* (Gatenby et al., 1985; Tabita and Small, 1985) and tobacco chloroplasts (Lin et al., 2014; Long et al., 2018) without additional chaperones. On the other hand, plant Rubisco has strict assembly requirements that can only be met by closely related species (Whitney et al., 2015; Sharwood, 2017). Some red algae have been reported to have Rubisco with the highest *S*_C/O_ values, which could potentially improve plant photosynthesis, but their Rubisco failed to assemble into active enzyme in tobacco due to lack of compatible assembly factors (Whitney et al., 2001; Lin and Hanson, 2018).

Previous studies on maize photosynthesis mutants lacking Rubisco identified three essential chaperones for Rubisco assembly: BSD2 (bundle-sheath defective 2), Raf1 and Raf2 (Rubisco accumulation factors 1 and 2) (Brutnell et al., 1999; Feiz et al., 2012; Feiz et al., 2014). Further structural studies have revealed the mechanism of Raf1 and BSD2 as well as RbcX in assisting the dimerization of large subunits and subsequent formation of the L_8_ core before the formation of the final holoenzyme (Liu et al., 2010; Hauser et al., 2015; Aigner et al., 2017).

In a recent breakthrough for the assembly of plant Rubisco, Aigner and co-workers were able to produce functional Arabidopsis Rubisco in *E. coli* by co-expressing at least five additional proteins: two plant-specific chaperonin (Cpn60α and Cpn60β) and three chaperones (Raf1, Raf2 and BSD2) (Aigner et al., 2017). They also demonstrated the assembly of a small amount of tobacco Rubisco by replacing the Arabidopsis *raf1* with a tobacco homolog. Needless to say, this advance will greatly facilitate functional studies of plant Rubisco. Although Arabidopsis has been a great model plant in genetic studies, tobacco offers several advantages in Rubisco engineering since both nuclear and chloroplast genome transformations can be readily carried out in tobacco followed by subsequent field trials of the transgenic plants with modified Rubisco to access their photosynthetic performance when standard agricultural practices are employed. Thus, an *E. coli* expression system with improved assembly of tobacco Rubisco will be useful to identify enzymes with improved catalytic properties whose performance can be tested in the field.

In the current study, we modified the expression vectors used by Aigner and co-workers (Aigner et al., 2017) to co-express tobacco chaperonins and chaperones. We also surveyed the expression of different small subunits in tobacco and co-expressed each of them in *E. coli* with the single tobacco chloroplast-encoded large subunit. We then quantified the tobacco Rubisco produced in *E. coli* with each small subunit and measured their carboxylation kinetics. We also found that the Rubisco produced in *E. coli* with a recently identified small subunit present in the secretory glands of tobacco trichomes (Laterre et al., 2017) has a considerably different kinetic profile.

## Results

### Tobacco Rubisco was successfully expressed and assembled in *E. coli* using a modified two-vector system

The pET and pCDF expression vectors we used in this study are based on the system recently reported by (Aigner et al., 2017). We obtained p*At*C60αβ/C20 and p*Nt*R1/*At*R2/B2 that were used in that study to co-express three Arabidopsis chaperonins (Cpn60α1, Cpn60β1 and Cpn20), two Arabidopsis chaperones (Raf2 and BSD2) and one tobacco chaperone (Raf1). Instead of expressing the two tobacco Rubisco subunits from a third vector, we introduced the tobacco *rbcL* and *rbcS-S1* genes into p*At*C60αβ/C20 and p*Nt*R1/*At*R2/B2 vectors along with arabinose-inducible P_BAD_ promoters to obtain pET-*Nt*L-*At*C60αβ20 and pCDF-*Nt*R1*At*R2B-*Nt*S1 respectively (Figure 1). We also introduced the tobacco *rbcX* gene into the latter since deleting *At-rbcX* was shown to reduce the yield of Arabidopsis Rubisco by up to 60% (Aigner et al., 2017).

**Figure 1.**
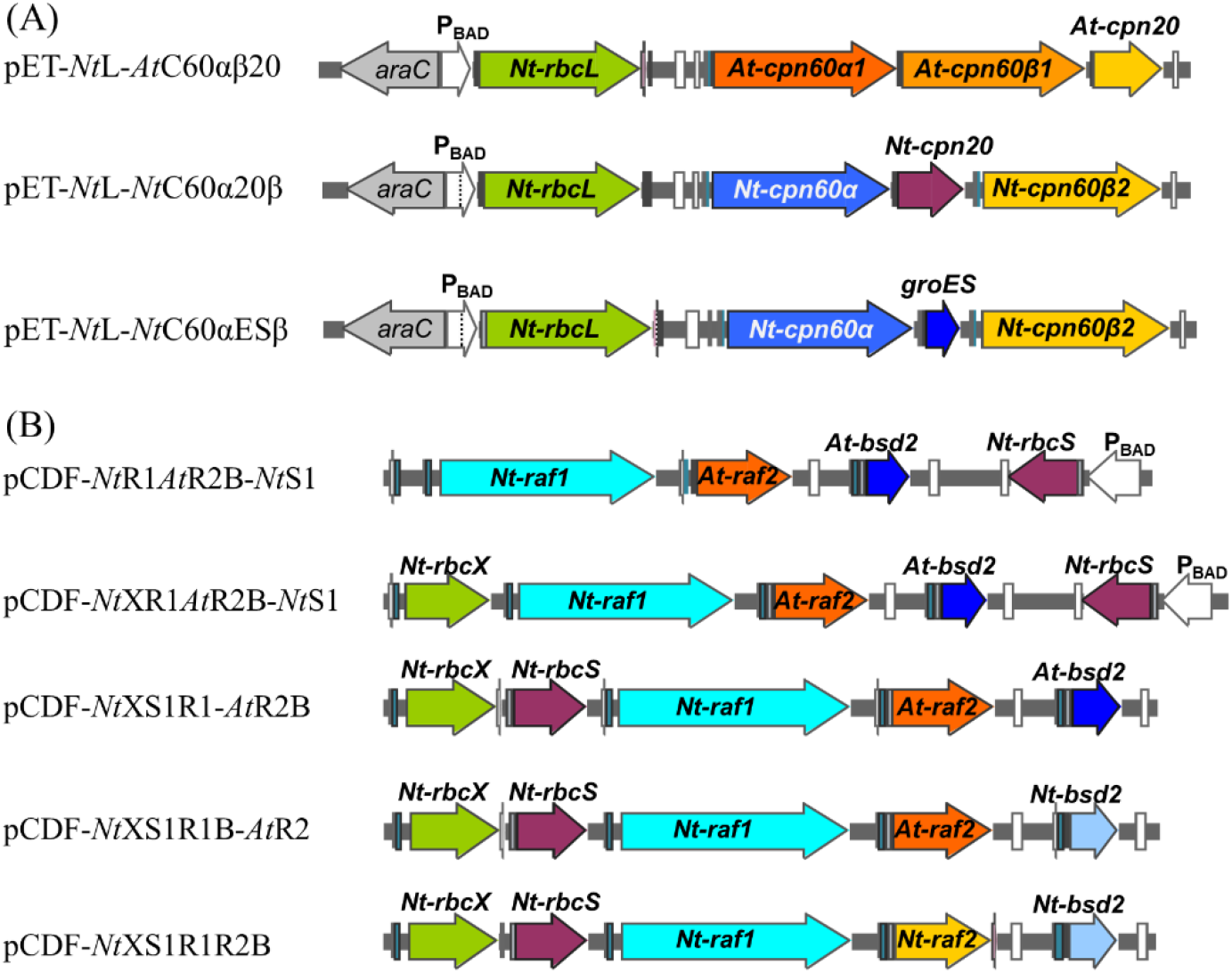
pET and pCDF *E. coli* expression vectors created in this study. (A) pET vectors express *Nt-rbcL* from P_BAD_ and *cpn60α*, *cpn60β* and either *cpn20* or *groES* from P_T7_. (B) pCDF vectors express *Nt-rbcS* from either P_BAD_ or P_T7_ and *rbcX*, *raf1*, *raf2* and *bsd2* from P_T7_.

Our modified two-vector system is compatible with *E. coli* expression strains that have extra tRNAs for rare *E. coli* codons such as Rosetta (DE3). We then followed the previously described growth protocol (Aigner et al., 2017) to initially induce the chaperonins and chaperones with IPTG for 3 hours at room temperature followed by the transfer of the *E. coli* cultures to a medium containing arabinose to induce the expression of tobacco Rubisco subunits for additional 16-18 hours at room temperature. We then analyzed the soluble extracts from three different *E. coli* strains on a native PAGE immunoblot with an antibody against Rubisco. We observed an L_8_S_8_ Rubisco complex band from the BL21(DE3) extract when the tobacco subunits were co-expressed with Arabidopsis chaperonins, Raf2, BSD2 and tobacco Raf1 (Figure 2). We also found that the Rosetta (DE3) strain was able to produce more L_8_S_8_ Rubisco than the two BL21(DE3) strains tested (Figure 3). Since none of the nine genes that were expressed in *E. coli* are codon-optimized for the bacterium, the presence of tRNAs for rare codons in the Rosetta strain likely resulted in improved expression.

**Figure 2.**
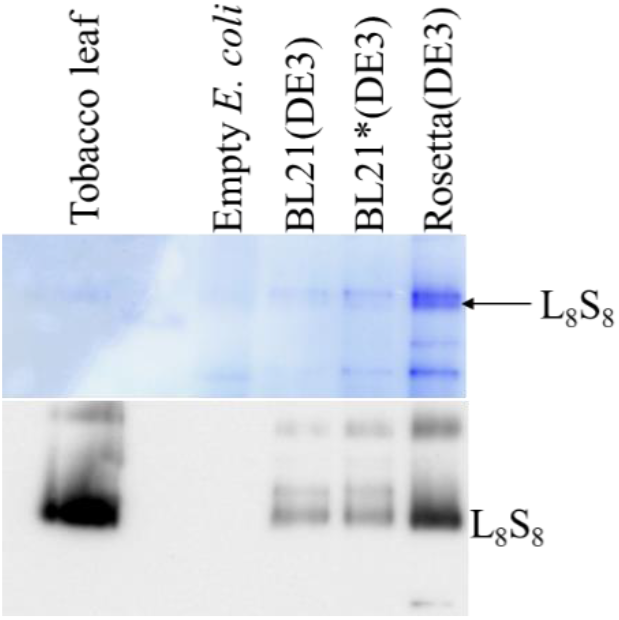
The native PAGE analysis of *E. coli* soluble extracts expressing genes from pET-*Nt*L-*At*C60αβ20 and pCDF-*Nt*XR1*At*R2B-*Nt*S1 with the top panel showing Coomassie blue staining and the bottom panel showing the immunoblot with the antibody against Rubisco. The genes expressed are *Nt-rbcL* (P_BAD_), *Nt-rbcS-S1* (P_BAD_), *Nt-rbcX*, *Nt-raf1*, *At-raf2*, *At-bsd2*, *At-cpn60α1*, *At-cpn60β1* and *At-cpn20*.

**Figure 3.**
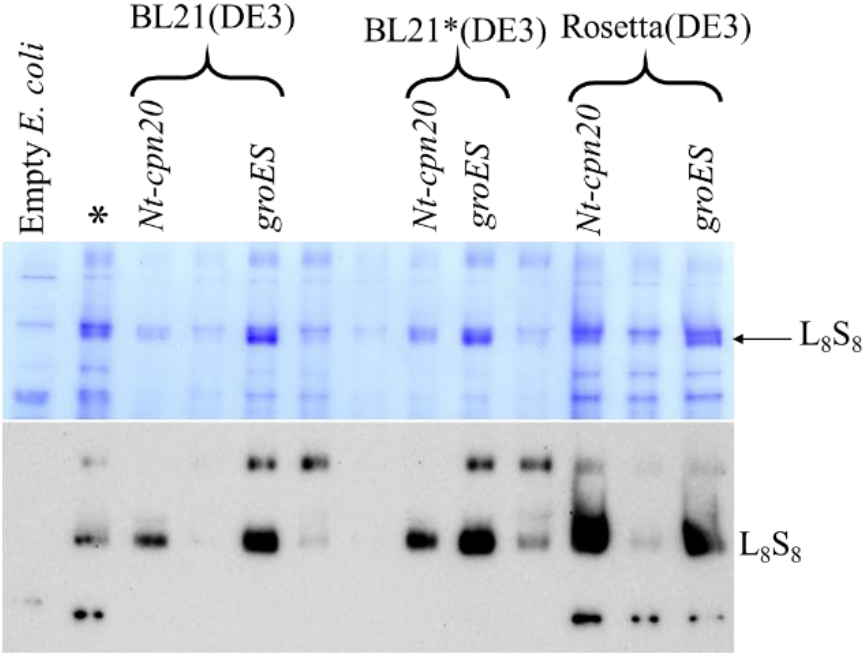
The native PAGE analysis of *E. coli* soluble extracts expressing genes from either pET-*Nt*L-*At*C60α20β or pET-*Nt*L-*Nt*C60αESβ and pCDF-*Nt*XS1R1-*At*R2B with the top panel showing Coomassie blue staining and the bottom panel showing the immunoblot with the antibody against Rubisco. The genes expressed are *Nt-rbcL* (P_BAD_), *Nt-rbcS-S1* (P_T7_), *Nt-rbcX*, *Nt-raf1*, *At-raf2*, *At-bsd2*, *Nt-cpn60α*, *Nt-cpn60β2* and either *Nt-cpn20* or *groES*. The lane marked with (*) was the Rosetta (DE3) extract expressing *At-cpn60α1*, *At-cpn60β2* and *At-cpn20*.

### The expression of tobacco Rubisco in *E. coli* was markedly improved when tobacco chaperonins were co-expressed

Next, we replaced the Arabidopsis chaperonins (Cpn60α1, Cpn60β1 and Cpn20) with tobacco counterparts. Alternatively, we also overexpressed *E. coli* cochaperonin GroES in place of tobacco Cpn20 since Aigner et al. (2017) showed that GroES was able to function with Arabidopsis Cpn60 to assemble Rubisco. We also tested the overexpression of tobacco RbcS with a T7 promoter (P_T7_) because the availability of RbcS was often limited in the assembly Arabidopsis Rubisco holoenzyme in *E. coli* (Aigner et al., 2017). All these changes resulted in dramatic improvements in tobacco Rubisco assembly in all three *E. coli* expression strains, especially when GroES was also overexpressed (Figure 3).

Subsequently, we replaced Arabidopsis *raf2* and *bsd2* genes in pCDF vectors with the tobacco versions and tested the expression of tobacco Rubisco in the BL21(DE3) strain. When the tobacco BSD2 was co-expressed instead of Arabidopsis BSD2, we observed increase in Rubisco expression although the improvement was not as dramatic as replacing Arabidopsis chaperonins with the tobacco ones (Figure 4). When Arabidopsis Raf2 was replaced with tobacco Raf2, there was no improvement in the assembly of Rubisco in *E. coli* (Figure 4). Although there appears to be no species specificity associated with Raf2 for Rubisco assembly, Raf2 is still important and necessary since the samples with lower tobacco Raf2 produced little Rubisco in *E. coli* (lanes marked with * in Figure 4).

**Figure 4.**
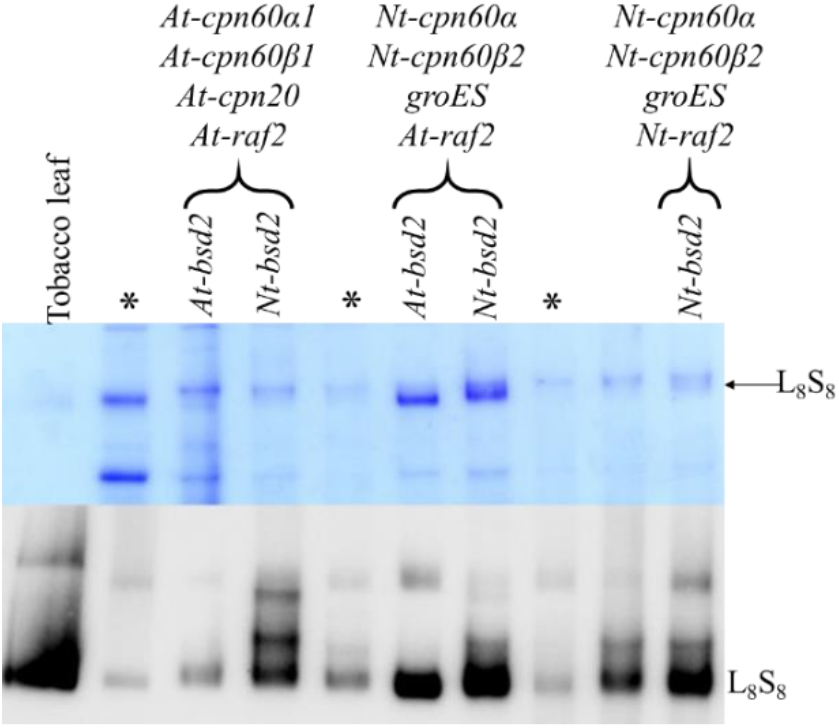
The native PAGE analysis of BL21 (DE3) *E. coli* soluble extracts expressing genes from either pET-*Nt*L-*At*C60αβ20 or pET-*Nt*L-*Nt*C60αESβ and either pCDF-*Nt*XS1R1-*At*R2B, pCDF-*Nt*XS1R1B-*At*R2 or pCDF-*Nt*XS1R1R2B with the top panel showing Coomassie blue staining and the bottom panel showing the immunoblot with the antibody against Rubisco. The genes expressed are *Nt-rbcL* (P_BAD_), *Nt-rbcS-S1* (P_T7_), *Nt-rbcX*, *Nt-raf1* and the other five genes as indicated on top of the lanes. The lanes marked with (*) were from *E. coli* strains expressing significantly less tobacco *raf2* due to difference in sequence near its start codon.

### Functional Rubisco can be produced in *E. coli* expressing different tobacco small subunits

In higher plants, the Rubisco small subunits are expressed from multiple nuclear genes, usually encoding slightly different amino acid sequences. In tobacco, 13 small subunit genes were previously identified (Gong et al., 2014). We analyzed their expression levels using publicly available RNA-Seq data generated from tobacco leaf samples (Sierro et al., 2014) and found that 11 of them are being transcribed, with *Nt-rbcS-T1* having the highest levels of transcripts for both young and mature leaves and *Nt-rbcS-S1a*, *Nt-rbcS-S1b*, *Nt-rbcS-S2* and *Nt-rbcS-S5* making up most of the remaining transcripts (Figure 5A). Several of these genes encode identical mature small subunits, resulting in 7 unique small subunits (SSU-S1, SSU-S2, SSU-S5, SSU-T1, SSU-T2, SSU-T4a and SSU-T5) (Figure 5B).

**Figure 5.**
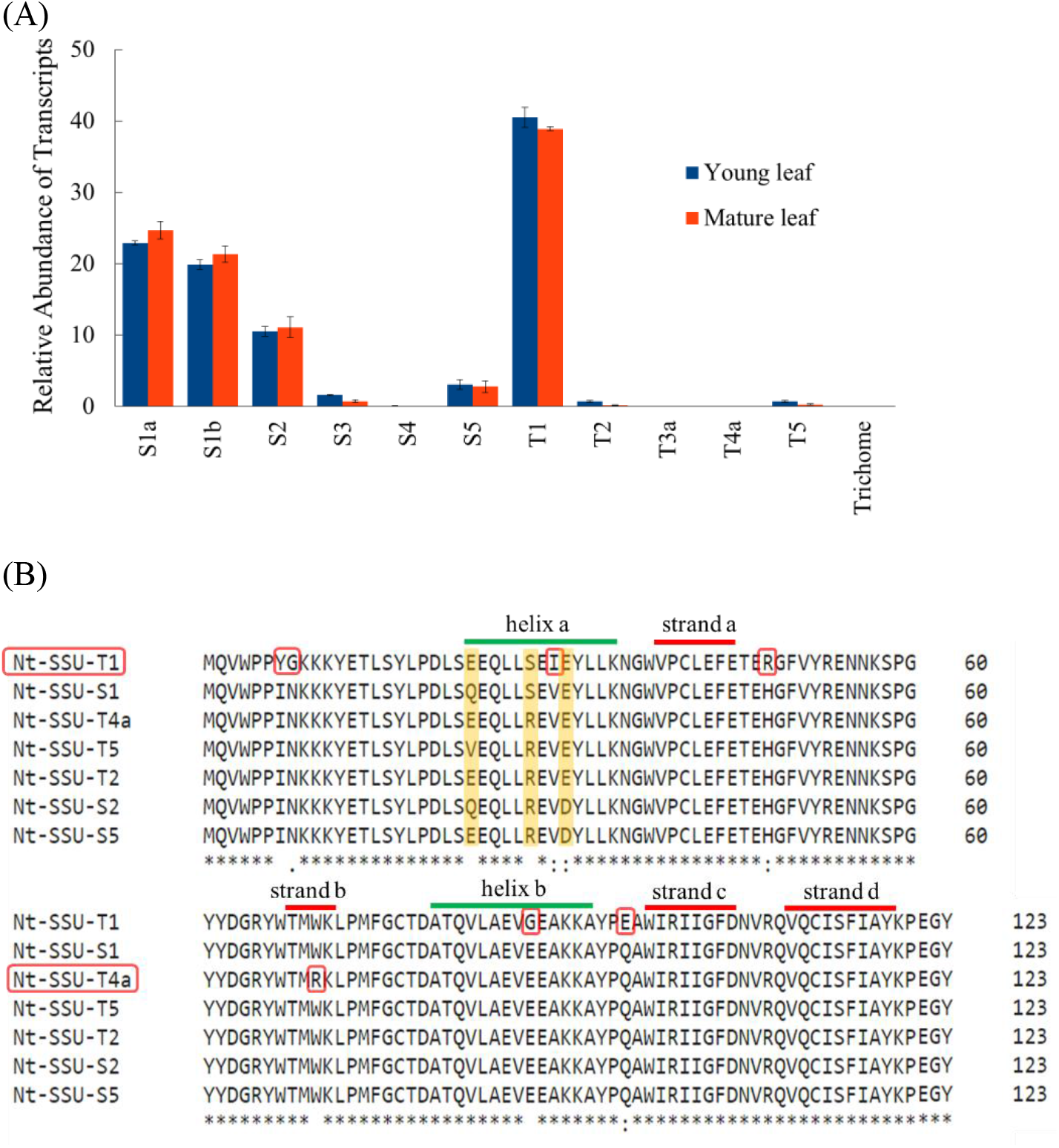
Survey of the Rubisco small subunits in tobacco. (A) Comparison of transcript abundances estimated from an RNA-Seq experiment of tobacco leaf tissue (Sierro et al., 2014) with Kallisto software (Bray et al., 2016). The error bars are standard deviations of the abundances obtained from three SRA files. (B) Multiple sequence alignment of seven unique tobacco small subunits with Clustal Omega. The chloroplast transit peptides are not included. The three variable residues in helix a are highlighted in yellow. The unique residues found only in Nt-SSU-T1 and Nt-SSU-T4a are indicated with red rectangles.

We co-expressed the tobacco *rbcL* and each of these 7 small subunits in BL21(DE3) together with two tobacco chaperonins (Cpn60α, Cpn60β), four tobacco chaperones (Raf1, Raf2, BSD2, RbcX) and *E. coli* GroES. We were able to detect an L_8_S_8_ Rubisco band similar to that from a tobacco leaf extract on a native PAGE immunoblot for 5 out of 7 small subunits. For *E. coli* co-expressing either *Nt-rbcS-T1* or *Nt-rbcS-T4a*, a slower-migrating band was instead observed (Figure 6). Next, we used these *E. coli* extracts to measure their RuBP carboxylation activities using ^14^C bicarbonate in the absence of O_2_. We found that the *E. coli* soluble extract with each of the five small subunits that produced L_8_S_8_ Rubisco bands displayed RuBP carboxylation activities (Figure 6). Surprisingly, of the two *E. coli* extracts that did not produce a normal L_8_S_8_ Rubisco band on the native PAGE immunoblot, the one co-expressing SSU-T1 was also able to carboxylate RuBP (Figure 6). We then quantified the Rubisco active sites in each of these extracts using ^14^C-CABP binding assays to determine the yield of Rubisco from these *E. coli* cultures (Kubien et al., 2011). We found that the *E. coli* strain expressing SSU-T1 produced higher levels of Rubisco active sites than the strains expressing four other SSUs (Figure 7).

**Figure 6.**
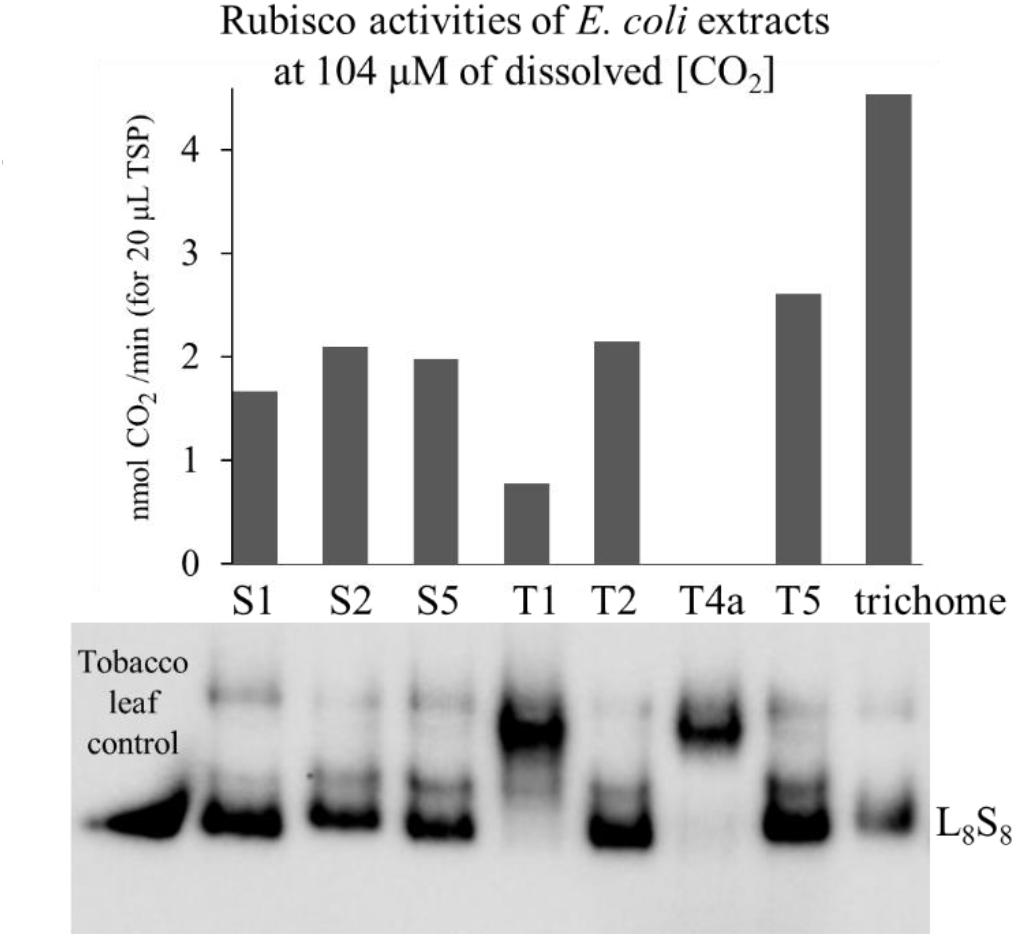
Comparison of tobacco Rubisco expressed in BL21 (DE3) with different small subunits. RuBP carboxylation rates shown on the top are averages of two measurements each using ^14^C bicarbonate. The native PAGE immunoblot in the bottom was obtained with the antibody against Rubisco.

**Figure 7.**
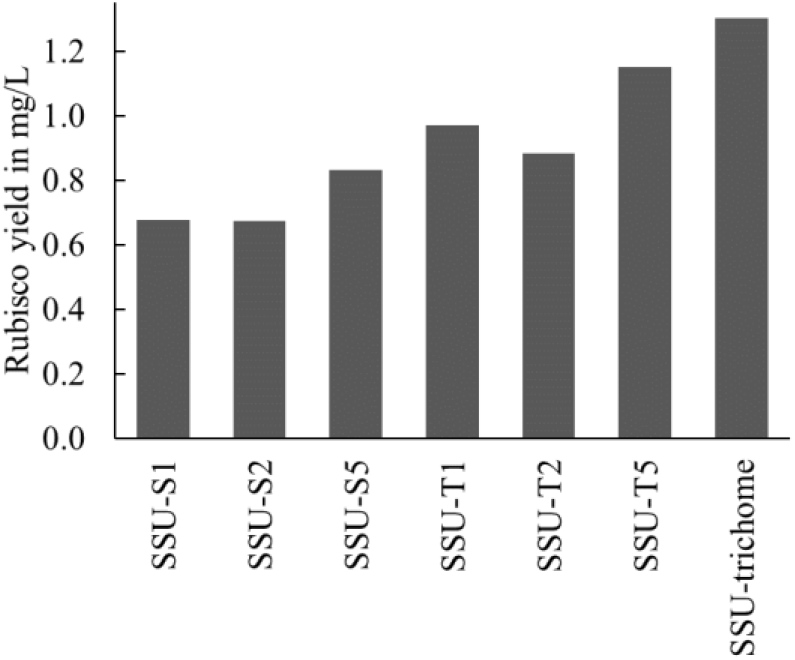
The yields of tobacco Rubisco from BL21 (DE3) *E. coli* expressing different small subunits as determined by ^14^C-CABP binding assays. The yields were based on 45 mL cultures. The values plotted are averages of two measurements each.

We also measured the RuBP carboxylation rates at six different dissolved CO_2_ levels for each *E. coli* extract and fitted the data to the standard Michaelis-Menten model to estimate their *K_C_* and *k_cat_* values. We found that the Rubisco produced in *E. coli* with each of the five small subunits (S1, S2, S5, T2 and T5) has an overall carboxylation kinetic profile that is very similar to the native Rubisco from the tobacco leaf although the enzymes produced in *E. coli* have slightly higher *K_C_* values (Figure 8). In contrast, the enzyme produced in *E. coli* with SSU-T1 displayed an inferior carboxylation activity due to its higher *K_C_* and lower *k_cat_* (Figure 8) in comparison to native Rubisco.

**Figure 8.**
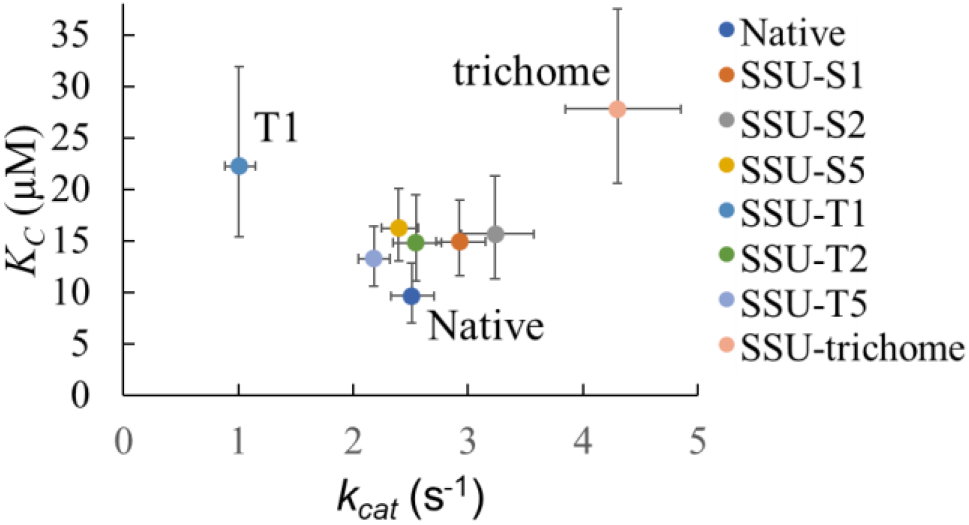
The kinetics of tobacco Rubisco expressed in *E. coli* with different small subunits compared to the native tobacco Rubisco in the absence of O_2_. The Michaelis-Menten constants for CO_2_ (*K_C_*) and turnover numbers (*k_cat_*) were obtained from nonlinear regression with the error bars showing 95% confidence intervals.

### The Rubisco enzyme assembled in *E. coli* with a phylogenetically diverse small subunit found in the tobacco trichome has significantly different kinetic properties

Recent studies discovered a phylogenetically diverse line of small subunits mainly expressed in nonphotosynthetic tissues of plants including tobacco trichomes (Morita et al., 2016; Laterre et al., 2017; Pottier et al., 2018). We co-expressed the trichome small subunit in BL21(DE3) with the same set of tobacco chaperonins and chaperones as described above. The corresponding *E. coli* extract had a normal L_8_S_8_ Rubisco band on the native PAGE immunoblot, and displayed the highest level of RuBP carboxylation activity as well as the highest level of Rubisco active sites (Figures 6 and 7). The tobacco trichome Rubisco produced in *E. coli* had carboxylation kinetics that are considerably different from the native enzyme found in mesophyll tissue and also different from those assembled in *E. coli* with other small subunits (Figure 8). Its higher turnover number and lower affinity for CO_2_ are consistent with the recent findings concerning tobacco trichome Rubisco (Laterre et al., 2017).

## Discussion

Aigner et al. (2017) recently overcame a major obstacle in functional studies of plant Rubisco by successfully expressing Arabidopsis Rubisco in *E. coli*. We modified two of the expression vectors used in their studies to enhance the expression of tobacco Rubisco in *E. coli*. Our two-vector system is compatible with *E. coli* expression systems with additional tRNAs for rare codons, which could be advantageous especially because none of the nine genes that are being co-expressed has codon usage optimized for *E. coli*. The ability to express plant Rubisco in *E. coli* means that both large and small subunits of this important enzyme can be rapidly modified for functional studies. Instead of expressing the two Rubisco subunits from a single vector, we expressed them from two different vectors in our two-vector system, which considerably reduces the number of vectors needed to express different combinations of large and small subunits.

We increased the assembly of tobacco Rubisco in *E. coli* by replacing Arabidopsis chaperonins and chaperones with tobacco versions. The most improvement was observed when the Arabidopsis Cpn60α, Cpn60β and Cpn20 with tobacco chaperonins Cpn60α, Cpn60β and the *E. coli* co-factor, GroES (Figure 4). In a previous report, co-expressing Arabidopsis *rbcL* and *raf1* in tobacco chloroplasts resulted in higher Rubisco levels than the levels achieved by expressing only Arabidopsis *rbcL* (Whitney et al., 2015). Similarly, tobacco BSD2 was able to provide additional improvement in tobacco Rubisco assembly in our current study. However, co-expressing tobacco Raf2 no longer improved the yield of tobacco Rubisco in *E. coli*. Raf2 was previously shown to be critical for Rubisco assembly in maize (Feiz et al., 2014) and Arabidopsis (Fristedt et al., 2018). Despite its importance, Arabidopsis Raf2 appeared to be fully functional in *E. coli* strains expressing tobacco Rubisco. Currently, Raf2 is the only Rubisco assembly chaperone, which mechanism has yet to be solved, and further structural studies of Raf2 in complex with Rubisco subunits will be critical to identify its role in Rubisco assembly.

Our current study shows that the RuBP carboxylation kinetics of tobacco Rubisco expressed in *E. coli* and the native enzyme from tobacco leaf are very similar but not identical (Figure 8). Clearly, the composition of small subunits in the native enzyme is heterogeneous, unlike the enzymes expressed in *E. coli*. Thus, it may be inappropriate to directly compare their kinetic properties. Interestingly, all five tobacco Rubisco, each with one of five different small subunits (S1, S2, S5, T2 and T5) expressed in *E. coli* have slightly higher *K_C_* values than the native enzyme, indicating that there may be some fundamental differences in Rubisco assembled in *E. coli* other than the composition of small subunits (Figure 8). Rubisco in higher plants is known to undergo multiple conserved posttranslational modifications, and their exact roles are not well understood (Houtz et al., 2008). The conserved N-terminal acetylation of its large subunit and possibly other posttranslational modifications were found to be missing in the Arabidopsis Rubisco expressed in *E. coli* (Aigner et al., 2017). We hypothesize that some of these posttranslational modifications may be one of the strategies that plants have evolved to increase the CO_2_ affinity of Rubisco as they adjusted to the declining global atmospheric CO_2_ level over the last 20 million years prior to the industrial revolution (Pearson and Palmer, 2000). Clearly, further genetic, proteomic and biochemical studies will be necessary to address this issue. Despite the minor differences between tobacco Rubisco in *E. coli* and in tobacco leaves, we believe the two enzymes are sufficiently similar to make future functional studies informative.

Two tobacco small subunits, SSU-T1 and SSU-T4a, have unique residue substitutions, which may explain why they were unable to assemble proper Rubisco holoenzymes in *E. coli* (Figures 5 and 6). The substitution of conserved tryptophan 70 with arginine in SSU-T4a must have proven to be fatal because SSU-T4a and SSU-T2 are otherwise identical. Since the transcript level of SSU-T4a is negligible, it is likely that the nonfunctional SSU-T4a is physiologically irrelevant. On the other hand, SSU-T1 possesses the highest transcript level and has six unique residue substitutions. Surprisingly, the *E. coli* extract with SSU-T1 displayed some RuBP carboxylation activity with inferior kinetics despite lacking properly assembled holoenzyme (Figure 8). It is possible that some specific posttranslational modifications that are missing in either Rubisco subunit produced in *E. coli* are necessary for SSU-T1 to assemble into fully functional Rubisco.

We also found that the tobacco trichome small subunit was able assemble into functional Rubisco in *E. coli* with higher *k_cat_* and *K_C_* values. This trichome small subunit belongs to a phylogenetically distinct line of small subunits named T-type with only about 65% sequence identity to the canonical small subunits recently named M-type and usually expressed in non-photosynthetic tissues (Morita et al., 2016; Pottier et al., 2018). Previous studies with a rice T-type homolog (Morita et al., 2014) as well as the same tobacco trichome subunit (Laterre et al., 2017) indicated that these Rubisco enzymes exhibit higher *k_cat_* and *K_C_* values than those in mesophyll tissue. Although the active sites are located within dimers of large subunits in the holoenzyme, it has long been known that the small subunits can influence the carboxylation kinetics likely by modifying the conformation of the enzyme (Spreitzer, 2003). For example, studies in Chlamydomonas Rubisco indicated that changes along the interface between the large and small subunits resulted in an altered CO_2_/O_2_ specificity factor and other kinetic parameters (Spreitzer et al., 2005; Genkov and Spreitzer, 2009). In another study, a transgenic rice expressing a small subunit from sorghum, a C_4_ plant, had Rubisco with higher catalytic rates than the rice enzyme (Ishikawa et al., 2011).

Phylogenetic analysis of Rubisco has revealed that the substitution rates in the small subunits are much higher than those in the large subunit in higher plants (Andersson and Backlund, 2008). It is possible that diversification of the small subunits represents one strategy that plants have employed to adjust the kinetics of Rubisco as they adapt to different environmental conditions. Some information has been provided by analysis of *Flaveria* species, in which both C_3_ and C_4_ carbon fixation exists (Kapralov et al., 2011). Expression of variant and mutant small subunits in *E. coli* in the future should provide a wealth of information concerning their roles in Rubisco enzyme kinetics and assembly.

One potential approach to identify Rubisco with improved kinetics is directed evolution using Rubisco-dependent *E. coli* strains. This approach has been applied for Rubisco from cyanobacteria (Smith and Tabita, 2003; Mueller-Cajar and Whitney, 2008; Durao et al., 2015; Wilson et al., 2018), *Rhodospirillum rubrum* (Mueller-Cajar et al., 2007) and *Methanococcoides burtonii* (Wilson et al., 2016) and can be extended to plant Rubisco by co-expressing the required chaperonins and chaperones in *E. coli*. Recently, several studies measured detailed kinetics of Rubisco from a large number of plant species and identified potential kinetic switches in the large subunit (Galmés et al., 2014; Orr et al., 2016; Prins et al., 2016). Evolutionary analyses of the large subunits in C_4_ plants also proposed residues that could be important in controlling the kinetics of Rubisco (Christin et al., 2008; Kapralov et al., 2012; Studer et al., 2014). However, due to enormous effort required to modify Rubisco in plants, only a few studies actually examined the effects of altering Rubisco in plants (Whitney et al., 1999; Whitney et al., 2011b). Our current study demonstrates that extension of the *E. coli* system to Rubisco enzymes from important crops should be feasible by incorporating appropriate species-specific assembly factors.

## Materials and Methods

### 1. Construction of *E. coli* expression plasmids

We first cloned all the genes in Table 1 except for *Nt-rbcL* into BJFE holding vectors with a T7 promoter (P_T7_), a ribosome binding site (RBS) and restriction sites (AscI, NotI) for rapid transfer into expression vectors. All the genes expressed in this study are summarized in Table 1, and the *E. coli* expression vectors in Table 2. The information on oligonucleotides used in generation of these vectors and full sequences of all vectors will be available upon request or at the NCBI website once the manuscript is published in a peer-reviewed journal.

**Table 1.**
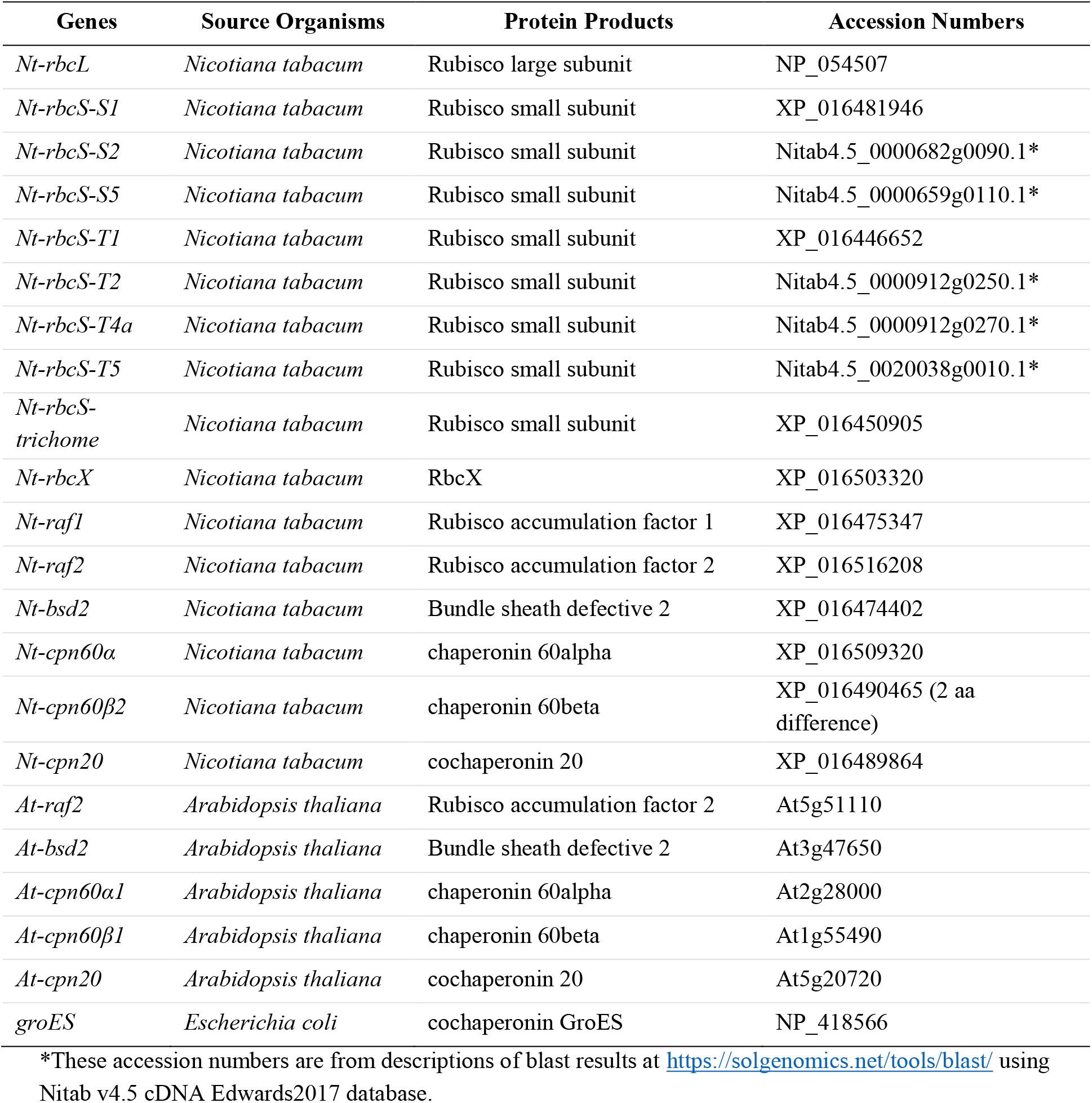
Information on the genes expressed in this study.

| Genes | Source Organisms | Protein Products | Accession Numbers |
| --- | --- | --- | --- |
| *Nt-rbcL* | *Nicotiana tabacum* | Rubisco large subunit | NP_054507 |
| *Nt-rbcS-S1* | *Nicotiana tabacum* | Rubisco small subunit | XP_016481946 |
| *Nt-rbcS-S2* | *Nicotiana tabacum* | Rubisco small subunit | Nitab4.5_0000682g0090.1* |
| *Nt-rbcS-S5* | *Nicotiana tabacum* | Rubisco small subunit | Nitab4.5_0000659g0110.1* |
| *Nt-rbcS-T1* | *Nicotiana tabacum* | Rubisco small subunit | XP_016446652 |
| *Nt-rbcS-T2* | *Nicotiana tabacum* | Rubisco small subunit | Nitab4.5_0000912g0250.1* |
| *Nt-rbcS-T4a* | *Nicotiana tabacum* | Rubisco small subunit | Nitab4.5_0000912g0270.1* |
| *Nt-rbcS-T5* | *Nicotiana tabacum* | Rubisco small subunit | Nitab4.5_0020038g0010.1* |
| *Nt-rbcS-trichome* | *Nicotiana tabacum* | Rubisco small subunit | XP_016450905 |
| *Nt-rbcX* | *Nicotiana tabacum* | RbcX | XP_016503320 |
| *Nt-raf1* | *Nicotiana tabacum* | Rubisco accumulation factor 1 | XP_016475347 |
| *Nt-raf2* | *Nicotiana tabacum* | Rubisco accumulation factor 2 | XP_016516208 |
| *Nt-bsd2* | *Nicotiana tabacum* | Bundle sheath defective 2 | XP_016474402 |
| *Nt-cpn60α* | *Nicotiana tabacum* | chaperonin 60α | XP_016509320 |
| *Nt-cpn60β2* | *Nicotiana tabacum* | chaperonin 60β | XP_016490465 (2 aa difference) |
| *Nt-cpn20* | *Nicotiana tabacum* | cochaperonin 20 | XP_016489864 |
| *At-raf2* | *Arabidopsis thaliana* | Rubisco accumulation factor 2 | At5g51110 |
| *At-bsd2* | *Arabidopsis thaliana* | Bundle sheath defective 2 | At3g47650 |
| *At-cpn60α1* | *Arabidopsis thaliana* | chaperonin 60α | At2g28000 |
| *At-cpn60β1* | *Arabidopsis thaliana* | chaperonin 60β | At1g55490 |
| *At-cpn20* | *Arabidopsis thaliana* | cochaperonin 20 | At5g20720 |
| *groES* | *Escherichia coli* | cochaperonin GroES | NP_418566 |

\*These accession numbers are from descriptions of blast results at https://solgenomics.net/tools/blast/ using Nitab v4.5 cDNA Edwards2017 database.

**Table 2.**
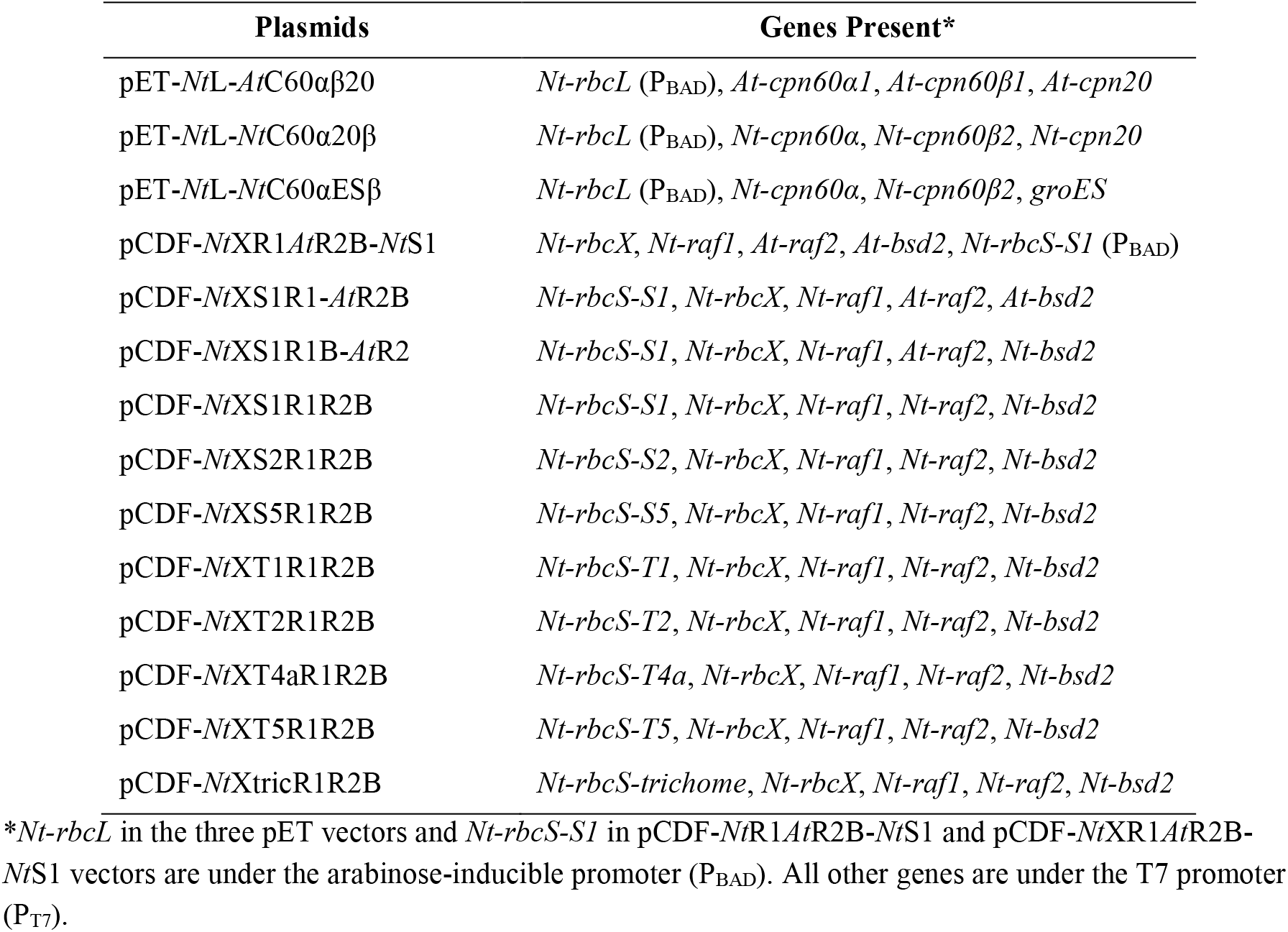
Summary of plasmids used in the expression of tobacco Rubisco in *E. coli*. Only the final plasmids used in the *E. coli* expression strains are included here.

| Plasmids | Genes Present* |
| --- | --- |
| pET-*NtL-AtC60αβ20* | *Nt-rbcL* (P<sub>BAD</sub>), *At-cpn60α1*, *At-cpn60β1*, *At-cpn20* |
| pET-*NtL-NtC60α20β* | *Nt-rbcL* (P<sub>BAD</sub>), *Nt-cpn60α*, *Nt-cpn60β2*, *Nt-cpn20* |
| pET-*NtL-NtC60αESβ* | *Nt-rbcL* (P<sub>BAD</sub>), *Nt-cpn60α*, *Nt-cpn60β2*, *groES* |
| pCDF-*NtXR1AtR2B-NtS1* | *Nt-rbcX*, *Nt-raf1*, *At-raf2*, *At-bsd2*, *Nt-rbcS-S1* (P<sub>BAD</sub>) |
| pCDF-*NtXS1R1-AtR2B* | *Nt-rbcS-S1*, *Nt-rbcX*, *Nt-raf1*, *At-raf2*, *At-bsd2* |
| pCDF-*NtXS1R1B-AtR2* | *Nt-rbcS-S1*, *Nt-rbcX*, *Nt-raf1*, *At-raf2*, *Nt-bsd2* |
| pCDF-*NtXS1R1R2B* | *Nt-rbcS-S1*, *Nt-rbcX*, *Nt-raf1*, *Nt-raf2*, *Nt-bsd2* |
| pCDF-*NtXS2R1R2B* | *Nt-rbcS-S2*, *Nt-rbcX*, *Nt-raf1*, *Nt-raf2*, *Nt-bsd2* |
| pCDF-*NtXS5R1R2B* | *Nt-rbcS-S5*, *Nt-rbcX*, *Nt-raf1*, *Nt-raf2*, *Nt-bsd2* |
| pCDF-*NtXT1R1R2B* | *Nt-rbcS-T1*, *Nt-rbcX*, *Nt-raf1*, *Nt-raf2*, *Nt-bsd2* |
| pCDF-*NtXT2R1R2B* | *Nt-rbcS-T2*, *Nt-rbcX*, *Nt-raf1*, *Nt-raf2*, *Nt-bsd2* |
| pCDF-*NtXT4aR1R2B* | *Nt-rbcS-T4a*, *Nt-rbcX*, *Nt-raf1*, *Nt-raf2*, *Nt-bsd2* |
| pCDF-*NtXT5R1R2B* | *Nt-rbcS-T5*, *Nt-rbcX*, *Nt-raf1*, *Nt-raf2*, *Nt-bsd2* |
| pCDF-*NtXtrichR1R2B* | *Nt-rbcS-trichome*, *Nt-rbcX*, *Nt-raf1*, *Nt-raf2*, *Nt-bsd2* |

\**Nt-rbcL* in the three pET vectors and *Nt-rbcS-S1* in pCDF-*NtR1AtR2B-NtS1* and pCDF-*NtXR1AtR2B-NtS1* vectors are under the arabinose-inducible promoter (P<sub>BAD</sub>). All other genes are under the T7 promoter (P<sub>T7</sub>).

(i) pET-*Nt*L-*At*C60αβ20 – *NtrbcL-S2A* which has alanine in place of serine at the second residue was amplified from tobacco DNA and ligated into NcoI-HindIII sites of pBAD-Dest49. Site-directed mutagenesis was next carried out to convert *NtrbcL-S2A* back into the wild-type sequence to obtain pBAD-*Nt*-rbcL vector. Next, *araC-P_BAD_-Nt-rbcL-term* was amplified from the vector and ligated into NaeI sites of p*At*C60αβ/C20 (Aigner et al., 2017).
(ii) pET-*Nt*L-*Nt*C60α20β – The fragment between the two MluI sites in pET-*Nt*L-*At*C60αβ20 was removed to generate pET-*Nt*L-*At*C60αβ20-MluI vector. NheI-RBS-*Nt*cpn60α-P_T7_-RBS-*Nt*cpn60β2-BsiWI was amplified from pRSF-*Nt*20αβ vector and ligated into XbaI and Acc65I sites of pET-*Nt*L-*At*C60αβ20-MluI to generate pET-*Nt*C60αβ-2MluI. Site-directed mutagenesis was then carried out to remove the MluI site in *Nt*cpn60β2 to generate pET-*Nt*C60αβ-1MluI. The fragment between the two MluI sites in pET-*Nt*L-*At*C60αβ20 was then inserted into pET-*Nt*C60αβ-1MluI to obtain pET-*Nt*L-*Nt*C60αβ. MauBI-NotI fragment from BJFE-RBS-*Nt*cpn20 was then ligated into AscI-NotI sites of pET-*Nt*L-*Nt*C60αβ to obtain pET-*Nt*L-*Nt*C60α20β.
(iii) pET-*Nt*L-*Nt*C60αESβ – MauBI-NotI fragment from BJFE-P_T7_-RBS-groES was ligated into AscI-NotI sites of pET-*Nt*L-*Nt*C60αβ to generate pET-*Nt*L-*Nt*C60αESβ.
(iv) pCDF-*Nt*XR1*At*R2B-*Nt*S1 – pCDF-*Nt*R1*At*R2B was obtained from Manajit Hayer-Hartl at Max Planck Institute of Biochemistry, Martinsried, Germany. Kpn2I-P_BAD_-*Nt*S1-B1002-Kpn2I was generated with overlapping PCR and ligated into the AgeI site of pCDF-*Nt*R1*At*R2B to obtain pCDF-*Nt*R1*At*R2B-*Nt*S1. NcoI-*Nt*rbcX-NotI was then ligated into pCDF-*Nt*R1*At*R2B-*Nt*S1 to obtain pCDF-*Nt*XR1*At*R2B-*Nt*S1.
(v) pCDF-*Nt*XS1R1-*At*R2B – NcoI-*Nt*rbcX-NotI was ligated into pCDF-*Nt*R1*At*R2B to obtain pCDF-*Nt*XR1*At*R2B. MauBI-NotI fragment from BJFE-P_T7_-RBS-*Nt*S1 was then ligated into AscI-NotI sites of pCDF-*Nt*XR1*At*R2B to obtain pCDF-*Nt*XS1R1-*At*R2B.
(vi) pCDF-*Nt*XS1R1B-*At*R2 – IG2-NtB-IG3 was generated with overlapping PCR and ligated into SpeI-XhoI sites of pCDF-*Nt*XS1R1-*At*R2B to obtain pCDF-*Nt*XS1R1B-*At*R2.
(vii) pCDF-*Nt*XS1R1R2B – IG1-NtR2-IG2 was generated with overlapping PCR and ligated into PstI-SpeI sites of pCDF-*Nt*XS1R1B-*At*R2 to obtain pCDF-*Nt*XS1R1R2B.
(viii) pCDF-*Nt*X(SSU)R1R2B – *Nt*R2B from pCDF-*Nt*XS1R1R2B was ligated into PstI-XhoI sites of pCDF-*Nt*XR1*At*R2B to obtain pCDF-*Nt*XR1R2B. Each MauBI-NotI fragment from BJFE-P_T7_-RBS-*Nt*SSU-(S2/S5/T1/T2/T4a/T5/trichome) was ligated into AscI, NotI sites of pCDF-*Nt*XR1R2B to obtain pCDF-*Nt*X(SSU)R1R2B with different tobacco small subunit genes after the *Nt-rbcX* gene.

### 2. Expression of tobacco Rubisco in *E. coli*

The *E. coli* cultures were grown at 37 °C 250 rpm overnight in LB medium and diluted 60-200 times into 6 mL or 45 mL LB. Ampicillin, spectinomycin and chloramphenicol were added at 150, 100 and 30 µg mL^−1^ respectively into the growth medium as necessary. Once the OD_600_ reached at about 0.3, the cultures were induced with 300 µM IPTG and grown at room temperature 250 rpm for 3 hours. They were then pelleted at 2500 rcf at room temperature for 5 minutes, resuspended in fresh LB medium containing 0.4% arabinose and grown at room temperature 250 rpm for 16-18 hours. They were then pelleted again and stored at −80 °C.

### 3. Native PAGE of *E. coli* extracts

The *E. coli* pellets were resuspended in 0.5 – 1.0 mL of lysis buffer consisting of 50 mM Tris-HCl pH 8, 10 mM MgCl_2_, 1 mM EDTA, 2 mM DTT and protease inhibitor cocktail (Thermo Scientific) and broken by sonication on ice. For tobacco leaf sample, the leaf tissues were ground in a Wheaton homogenizer in 100 mM Bicine/NaOH pH 8.0, 10 mM MgCl_2_, 1 mM EDTA, 1 mM ε–aminocaproic acid, 1 mM benzamidine, protease inhibitor cocktail, 1 mM phenylmethanesulfonyl fluoride, 1 mM KH_2_PO_4_, 2% w/v poly(ethylene glycol) 4000, 10 mM NaHCO_3_ and 10 mM DTT. After the cell debris were removed by centrifugation at 16,000 rcf for 5 minutes, the protein concentrations in the supernatants were estimated by the Bradford assay. About 20 µg of total soluble proteins of each sample was loaded to 3-12% Bis-Tris 1.0 mm Invitrogen NativePAGE^TM^ protein gels from ThermoFisher Scientific. Invitrogen NativePAGE^TM^ running buffer from ThermoFisher Scientific was used as the running buffer with 0.002% Coomassie Brilliant Blue G-250 added to the cathode buffer. The native PAGE was run at 4 °C at 150-250 V for about 1.5 hours and then transferred to PVDF membranes with 0.45 um removal rating at 100 V for 1 hour. The membranes were then blocked with 5% milk in TBST buffer at room temperature for 1 hour and incubated with the antibody against Rubisco (kindly provided by P. John Andralojc from Rothamsted Research) in 5% milk in TBST buffer at 4 °C overnight. The primary antibody was detected with an HRP secondary antibody in 2.5% milk in TBST buffer at room temperature and chemiluminescence was recorded with the ChemiDoc^TM^ MP Imaging System from Bio-Rad.

### 4. Quantification of relative expression of *rbcS* genes in Tobacco

Publicly available SRA files used for tobacco genome sequencing (NCBI Bioproject-PRJNA208209) (Sierro et al., 2014) were utilized to quantitate relative abundance of genes. Three transcriptomic SRA files each for young leaf and mature leaf were used to quantitate relative abundance of *rbcS* genes using Kallisto (Bray et al., 2016) with standard parameters.

### 5. Quantification of Rubisco active sites using ^14^C-CABP

A previously described protocol (Whitney and Sharwood, 2014) was followed to synthesize ^14^C-CABP from RuBP (Sigma-Aldrich part number 83895) and ^14^C-KCN (PerkinElmer). 40-100 µL of each sample was incubated with 7.2 nmol ^14^C-CABP with 5 mCi/mmol of specific activity for at least 20 minutes and applied to size-exclusion chromatography with Sephadex G50 Fine (Santa Cruz Biotechnology) as described previously (Kubien et al., 2011). The eluted fractions were mixed with Ultima Gold liquid scintillation cocktail from PerkinElmer and the activities of Rubisco-bound ^14^C-CABP were counted with a Beckman LS 6000IC scintillation counter.

### 6. Determination of RuBP carboxylation kinetics

About 15 mL of assay buffer consisting of 110 mM Bicine-NaOH and 22 mM MgCl_2_ at pH 8.0 was equilibrated with CO_2_-free N_2_ gas for at least 1 hour and mixed with about 1 mg of carbonic anhydrase (Sigma-Aldrich). 915 µL was then added to individual two-dram glass vials, which were then sealed with open top caps with bonded PTFE faced silicone liners (Wheaton part number W240842), and equilibrated again with CO_2_-free N_2_ gas for at least another 30 minutes at 25 °C. Six different concentrations of 50 µL of ^14^C bicarbonate were then added the vials to achieve six final bicarbonate concentrations ranging from 10.55 mM to 177.82 mM. 15 µL of 26.7 mM RuBP (Santa Cruz Biotechnology, sc-214827A) was then added to each vial. The reaction was then initiated by the addition of 20 µL of each sample containing Rubisco to the vials and stopped exactly one minute later by the addition of 200 µL of 20% (v/v) formic acid. The caps were then removed from the vials, which were then left on a heating block set at about 100 °C. Once almost all solutions in the vials were evaporated, 0.5 mL of ddH_2_O was added to each vial to dissolve the leftover chemicals. Each vial was mixed with at least 3.5 mL of Ultima Gold liquid scintillation cocktail from PerkinElmer and the acid-stable ^14^C compounds in the vials were counted with a Beckman LS 6000IC scintillation counter. The catalytic rates at different [CO_2_] concentrations were then fitted to the standard Michaelis-Menten equation with nonlinear regression in R software to obtain the *K_C_*, *V_max_* and 95% confidence intervals. The value of *k_cat_* was then obtained by dividing *V_max_* with the Rubisco active sites in each sample.

## Acknowledgments

We thank Manajit Hayer-Hartl from Max Planck Institute of Biochemistry in Martinsried, Germany, for providing us with their *E. coli* expression vectors. We also thank Douglas Orr, Elizabete Carmo-Silva and Martin Parry from Lancaster University in Lancaster, UK for advice on experiments to quantify Rubisco active sites and measure RuBP carboxylation kinetics. This study is funded by the Chemical Sciences, Geosciences, and Biosciences Division, Office of Basic Energy Sciences, Office of Science, U.S. Department of Energy to M.R.H. (Award Number: DE-SC0014339).

## References

1. Aigner, H., Wilson, R.H., Bracher, A., Calisse, L., Bhat, J.Y., Hartl, F.U., and Hayer-Hartl, M. (2017). Plant RuBisCo assembly in *E. coli* with five chloroplast chaperones including BSD2. Science 358, 1272–1278.

2. Andersson, I., and Backlund, A. (2008). Structure and function of Rubisco. Plant Physiol. Biochem. 46, 275–291.

3. Bauwe, H., Hagemann, M., Kern, R., and Timm, S. (2012). Photorespiration has a dual origin and manifold links to central metabolism. Curr. Opin. Plant Biol. 15, 269–275.

4. Bray, N.L., Pimentel, H., Melsted, P., and Pachter, L. (2016). Near-optimal probabilistic RNA-seq quantification. Nat. Biotechnol. 34, 525–527.

5. Brutnell, T.P., Sawers, R.J., Mant, A., and Langdale, J.A. (1999). BUNDLE SHEATH DEFECTIVE2, a novel protein required for post-translational regulation of the *rbcL* gene of maize. Plant Cell 11, 849–864.

6. Christin, P.-A., Salamin, N., Muasya, A.M., Roalson, E.H., Russier, F., and Besnard, G. (2008). Evolutionary switch and genetic convergence on *rbcL* following the evolution of C_4_ photosynthesis. Mol. Biol. Evol. 25, 2361–2368.

7. Durao, P., Aigner, H., Nagy, P., Mueller-Cajar, O., Hartl, F.U., and Hayer-Hartl, M. (2015). Opposing effects of folding and assembly chaperones on evolvability of Rubisco. Nat. Chem. Biol. 11, 148–155.

8. Evans, J.R. (1989). Photosynthesis and nitrogen relationships in leaves of C_3_ plants. Oecologia 78, 9–19.

9. Feiz, L., Williams-Carrier, R., Wostrikoff, K., Belcher, S., Barkan, A., and Stern, D.B. (2012). Ribulose-1,5-bis-phosphate carboxylase/oxygenase accumulation factor1 is required for holoenzyme assembly in maize. Plant Cell 24, 3435–3446.

10. Feiz, L., Williams-Carrier, R., Belcher, S., Montano, M., Barkan, A., and Stern, D.B. (2014). A protein with an inactive pterin-4a-carbinolamine dehydratase domain is required for Rubisco biogenesis in plants. Plant J. 80, 862–869.

11. Fristedt, R., Hu, C., Wheatley, N., Roy, L.M., Wachter, R.M., Savage, L., Harbinson, J., Kramer, D.M., Merchant, S.S., Yeates, T., and Croce, R. (2018). RAF2 is a RuBisCO assembly factor in *Arabidopsis thaliana*. Plant J. 94, 146–156.

12. Galmés, J., Kapralov, M.V., Andralojc, P.J., Conesa, M.À., Keys, A.J., Parry, M.A.J., and Flexas, J. (2014). Expanding knowledge of the Rubisco kinetics variability in plant species: environmental and evolutionary trends. Plant Cell Environ. 37, 1989–2001.

13. Gatenby, A.A., van der Vies, S.M., and Bradley, D. (1985). Assembly in *E. coli* of a functional multi-subunit ribulose bisphosphate carboxylase from a blue-green alga. Nature 314, 617–620.

14. Genkov, T., and Spreitzer, R.J. (2009). Highly conserved small subunit residues influence rubisco large subunit catalysis. J. Biol. Chem. 284, 30105–30112.

15. Gong, L., Olson, M., and Wendel, J.F. (2014). Cytonuclear evolution of rubisco in four allopolyploid lineages. Mol. Biol. Evol. 31, 2624–2636.

16. Hauser, T., Bhat, J.Y., Milicic, G., Wendler, P., Hartl, F.U., Bracher, A., and Hayer-Hartl, M. (2015). Structure and mechanism of the Rubisco-assembly chaperone Raf1. Nat. Struct. Mol. Biol. 22, 720–728.

17. Houtz, R.L., Magnani, R., Nayak, N.R., and Dirk, L.M. (2008). Co- and post-translational modifications in Rubisco: unanswered questions. J. Exp. Bot. 59, 1635–1645.

18. Ishikawa, C., Hatanaka, T., Misoo, S., Miyake, C., and Fukayama, H. (2011). Functional incorporation of sorghum small subunit increases the catalytic turnover rate of Rubisco in transgenic rice. Plant Physiol. 156, 1603–1611.

19. Jordan, D.B., and Ogren, W.L. (1981). A sensitive assay procedure for simultaneous determination of ribulose-1,5-bisphosphate carboxylase and oxygenase activities. Plant Physiol. 67, 237–245.

20. Kapralov, M.V., Smith, J.A.C., and Filatov, D.A. (2012). Rubisco evolution in C_4_ eudicots: an analysis of Amaranthaceae *Sensu Lato*. PLoS One 7, e52974.

21. Kapralov, M.V., Kubien, D.S., Andersson, I., and Filatov, D.A. (2011). Changes in Rubisco kinetics during the evolution of C_4_ photosynthesis in *Flaveria* (Asteraceae) are associated with positive selection on genes encoding the enzyme. Mol. Biol. Evol. 28, 1491–1503.

22. Kubien, D.S., Brown, C.M., and Kane, H.J. (2011). Quantifying the amount and activity of Rubisco in leaves. Methods Mol. Biol. 684, 349–362.

23. Laterre, R., Pottier, M., Remacle, C., and Boutry, M. (2017). Photosynthetic trichomes contain a specific Rubisco with a modified pH-dependent activity. Plant Physiol. 173, 2110–2120.

24. Lin, M.T., and Hanson, M.R. (2018). Red algal Rubisco fails to accumulate in transplastomic tobacco expressing *Griffithsia monilis RbcL* and *RbcS* genes. Plant Direct 2, e00045.

25. Lin, M.T., Occhialini, A., Andralojc, P.J., Parry, M.A., and Hanson, M.R. (2014). A faster Rubisco with potential to increase photosynthesis in crops. Nature 513, 547–550.

26. Liu, C., Young, A.L., Starling-Windhof, A., Bracher, A., Saschenbrecker, S., Rao, B.V., Rao, K.V., Berninghausen, O., Mielke, T., Hartl, F.U., Beckmann, R., and Hayer-Hartl, M. (2010). Coupled chaperone action in folding and assembly of hexadecameric Rubisco. Nature 463, 197–202.

27. Long, B.M., Hee, W.Y., Sharwood, R.E., Rae, B.D., Kaines, S., Lim, Y.L., Nguyen, N.D., Massey, B., Bala, S., von Caemmerer, S., Badger, M.R., and Price, G.D. (2018). Carboxysome encapsulation of the CO_2_-fixing enzyme Rubisco in tobacco chloroplasts. Nat. Commun. 9, 3570.

28. Morita, K., Hatanaka, T., Misoo, S., and Fukayama, H. (2014). Unusual small subunit that is not expressed in photosynthetic cells alters the catalytic properties of rubisco in rice. Plant Physiol. 164, 69–79.

29. Morita, K., Hatanaka, T., Misoo, S., and Fukayama, H. (2016). Identification and expression analysis of non-photosynthetic Rubisco small subunit, *OsRbcS1*-like genes in plants. Plant Gene 8, 26–31.

30. Mueller-Cajar, O., and Whitney, S.M. (2008). Evolving improved *Synechococcus* Rubisco functional expression in *Escherichia coli*. Biochem. J. 414, 205–214.

31. Mueller-Cajar, O., Morell, M., and Whitney, S.M. (2007). Directed evolution of rubisco in *Escherichia coli* reveals a specificity-determining hydrogen bond in the form II enzyme. Biochemistry 46, 14067–14074.

32. Orr, D.J., Alcantara, A., Kapralov, M.V., Andralojc, P.J., Carmo-Silva, E., and Parry, M.A. (2016). Surveying Rubisco diversity and temperature response to improve crop photosynthetic efficiency. Plant Physiol. 172, 707–717.

33. Pearson, P.N., and Palmer, M.R. (2000). Atmospheric carbon dioxide concentrations over the past 60 million years. Nature 406, 695–699.

34. Pottier, M., Gilis, D., and Boutry, M. (2018). The Hidden Face of Rubisco. Trends Plant Sci 23, 382–392.

35. Prins, A., Orr, D.J., Andralojc, P.J., Reynolds, M.P., Carmo-Silva, E., and Parry, M.A. (2016). Rubisco catalytic properties of wild and domesticated relatives provide scope for improving wheat photosynthesis. J. Exp. Bot. 67, 1827–1838.

36. Savir, Y., Noor, E., Milo, R., and Tlusty, T. (2010). Cross-species analysis traces adaptation of Rubisco toward optimality in a low-dimensional landscape. Proc. Natl. Acad. Sci. U. S. A. 107, 3475–3480.

37. Sharwood, R.E. (2017). Engineering chloroplasts to improve Rubisco catalysis: prospects for translating improvements into food and fiber crops. New Phytol. 213, 494–510.

38. Sierro, N., Battey, J.N.D., Ouadi, S., Bakaher, N., Bovet, L., Willig, A., Goepfert, S., Peitsch, M.C., and Ivanov, N.V. (2014). The tobacco genome sequence and its comparison with those of tomato and potato. Nat. Commun. 5, 3833.

39. Smith, S.A., and Tabita, F.R. (2003). Positive and negative selection of mutant forms of prokaryotic (cyanobacterial) ribulose-1,5-bisphosphate carboxylase/oxygenase. J. Mol. Biol. 331, 557–569.

40. Spreitzer, R.J. (2003). Role of the small subunit in ribulose-1,5-bisphosphate carboxylase/oxygenase. Arch. Biochem. Biophys. 414, 141–149.

41. Spreitzer, R.J., Peddi, S.R., and Satagopan, S. (2005). Phylogenetic engineering at an interface between large and small subunits imparts land-plant kinetic properties to algal Rubisco. Proc. Natl. Acad. Sci. U. S. A. 102, 17225–17230.

42. Studer, R.A., Christin, P.A., Williams, M.A., and Orengo, C.A. (2014). Stability-activity tradeoffs constrain the adaptive evolution of RubisCO. Proc. Natl. Acad. Sci. U. S. A. 111, 2223–2228.

43. Tabita, F.R., and Small, C.L. (1985). Expression and assembly of active cyanobacterial ribulose-1,5-bisphosphate carboxylase/oxygenase in *Escherichia coli* containing stoichiometric amounts of large and small subunits. Proc. Natl. Acad. Sci. U. S. A. 82, 6100–6103.

44. Tabita, F.R., Hanson, T.E., Satagopan, S., Witte, B.H., and Kreel, N.E. (2008). Phylogenetic and evolutionary relationships of RubisCO and the RubisCO-like proteins and the functional lessons provided by diverse molecular forms. Philos. Trans. R. Soc. Lond. B. Biol. Sci. 363, 2629–2640.

45. Tcherkez, G.G.B., Farquhar, G.D., and Andrews, T.J. (2006). Despite slow catalysis and confused substrate specificity, all ribulose bisphosphate carboxylases may be nearly perfectly optimized. Proc. Natl. Acad. Sci. U. S. A. 103, 7246–7251.

46. Walker, B.J., VanLoocke, A., Bernacchi, C.J., and Ort, D.R. (2016). The costs of photorespiration to food production now and in the future. Annu. Rev. Plant. Biol. 67, 107–129.

47. Whitney, S.M., and Sharwood, R.E. (2014). Plastid transformation for Rubisco engineering and protocols for assessing expression. Methods Mol. Biol. 1132, 245–262.

48. Whitney, S.M., Houtz, R.L., and Alonso, H. (2011a). Advancing our understanding and capacity to engineer nature’s CO_2_-sequestering enzyme, Rubisco. Plant Physiol. 155, 27–35.

49. Whitney, S.M., von Caemmerer, S., Hudson, G.S., and Andrews, T.J. (1999). Directed mutation of the Rubisco large subunit of tobacco influences photorespiration and growth. Plant Physiol. 121, 579–588.

50. Whitney, S.M., Baldet, P., Hudson, G.S., and Andrews, T.J. (2001). Form I Rubiscos from non-green algae are expressed abundantly but not assembled in tobacco chloroplasts. Plant J. 26, 535–547.

51. Whitney, S.M., Birch, R., Kelso, C., Beck, J.L., and Kapralov, M.V. (2015). Improving recombinant Rubisco biogenesis, plant photosynthesis and growth by coexpressing its ancillary RAF1 chaperone. Proc. Natl. Acad. Sci. U. S. A. 112, 3564–3569.

52. Whitney, S.M., Sharwood, R.E., Orr, D., White, S.J., Alonso, H., and Galmes, J. (2011b). Isoleucine 309 acts as a C_4_ catalytic switch that increases ribulose-1,5-bisphosphate carboxylase/oxygenase (rubisco) carboxylation rate in *Flaveria*. Proc. Natl. Acad. Sci. U. S. A. 108, 14688–14693.

53. Wilson, R.H., and Hayer-Hartl, M. (2018). Complex chaperone dependence of Rubisco biogenesis. Biochemistry 57, 3210–3216.

54. Wilson, R.H., Alonso, H., and Whitney, S.M. (2016). Evolving *Methanococcoides burtonii* archaeal Rubisco for improved photosynthesis and plant growth. Sci. Rep. 6, 22284.

55. Wilson, R.H., Martin-Avila, E., Conlan, C., and Whitney, S.M. (2018). An improved *Escherichia coli* screen for Rubisco identifies a protein–protein interface that can enhance CO_2_-fixation kinetics. J. Biol. Chem. 293, 18–27.

